# A 3D Soft X-Ray Structural Atlas of Multicellular Cable Bacteria

**DOI:** 10.64898/2026.09.16.752168

**Authors:** Anaísa Coelho, Aneesh Deshmukh, Valentina Loconte, Magdalene A. MacLean, Carolyn A. Larabell, Mohamed Y. El-Naggar

## Abstract

The multicellular filamentous cable bacteria perform centimeter-scale electron transport via a continuous periplasmic conductive network. Resolving how this electrical architecture is coordinated with intracellular organization requires an integrated approach coupling 3D ultrastructure with *in situ* biochemical composition. Here, we integrate soft X-ray tomography (SXT) with cryo-electron tomography, elemental mapping, and fluorescence microscopy to generate a volumetric 3D atlas of intact cable bacteria filaments. Whole-cell reconstructions reveal physiological heterogeneity and cell-cycle-dependent reorganization of intracellular vesicles and granules. X-ray absorption measurements resolve biochemical differences within the physically continuous conductive apparatus, where periplasmic conductive fibers possess higher biomolecular density than junction lamellae at cell junctions. Furthermore, cell elongation preceding division is associated with two high-density, ring-like nucleoid-associated domains, accompanied by asymmetric membrane constriction and localized FtsZ assemblies. These findings establish SXT as a powerful approach for studying multicellular bacterial organization and provide a multi-scale framework for understanding the conductive machinery of cable bacteria.

## INTRODUCTION

Cable bacteria are multicellular filamentous bacteria of the deltaproteobacterial family *Desulfobulbaceae* that are unique in the microbial world due to their ability to perform long-distance electron transport (LDET)^1^. Currently classified into two candidate genera, the marine *Candidatus* Electrothrix and the freshwater *Candidatus* Electronema ^2–4^, these organisms couple sulfide oxidation in anoxic sediment layers to oxygen reduction at the sediment–water interface, generating electrical currents spanning centimeter-scale distances ^1,5^. This conductivity is facilitated by a highly specialized, continuous periplasmic network of longitudinal fibers shared by thousands of cells within a single filament ^6–8^. How this coordination is achieved within intact multicellular filaments remains largely unexplored.

Over the past decade, ultrastructural studies have provided insight into the architecture of cable bacteria ^1,6,9,10^. These studies identified the structural basis of the conductive network, revealing the longitudinal periplasmic conductive fibers (PCFs) that are proposed to contain sulfur-coordinated nickel cofactors ^11^. At cell–cell junctions, cryo-electron tomography (cryo-ET) further resolved junction lamellae (JL), comprising the outer membrane, a putative surface layer, and a central core lamella sheet (CLS), the latter physically connecting adjacent conductive fibers across cells ^9^. Additional intracellular structures, including membrane-bound vesicles and polyphosphate (poly-P) granules, have also been described ^9,12–15^. However, characterizing the native volumetric distribution of these structures and their spatial organization within intact multicellular filaments remains challenging. Conventional electron microscopy provides ultrastructural details but typically requires chemical fixation, staining, or sectioning, which can obscure the native spatial context ^16–19^. Cryo-ET circumvents these artifacts to capture cells in near-native states; however, the inelastic scattering of electrons intrinsically limits the penetrable sample thickness. Consequently, visualizing thicker specimens often involves targeted thinning, such as focused ion beam milling, which inherently complicates whole-cell volumetric analyses ^20,21^. Conversely, fluorescence microscopy allows visualization of entire filaments but lacks the spatial resolution required to study fine intracellular organization ^22^. This resolution-volume gap is particularly limiting for the multicellular cable bacteria, where function depends on coordination across multiple cells ^23,24^. Resolving how intracellular and conductive structures are spatially integrated within intact filaments requires an imaging modality capable of capturing fully hydrated, multicellular segments at high resolution.

Soft X-ray Tomography (SXT) bridges this resolution-volume gap. Operating within the ‘water window’ (284 to 543 eV), SXT exploits the natural contrast between water and carbon- and nitrogen-rich molecules. SXT enables quantitative three-dimensional (3D) mapping of intact, cryo-preserved cells in a near-native state, circumventing the need for chemical fixation, staining, or physical sectioning ^25–28^. Because X-ray absorption follows the Beer–Lambert law, each voxel is associated with a quantitative linear absorption coefficient (LAC), allowing intracellular structures to be differentiated based on their biochemical composition ^29^. Since different biomolecules absorb X-rays distinctively, LAC measurements allow the identification of carbon-dense structures (such as lipid-rich vesicles, condensed nucleoids, and protein-rich regions) from mineral-rich inclusions (like poly-P granules) and the less dense aqueous cytoplasm *in situ* ^25,27,28^. Despite these advantages, SXT has seen limited application in microbiology, where most studies focused on unicellular model organisms, including *Escherichia coli* and *Pseudomonas*, or intracellular pathogens within host cells ^25,30^. Consequently, the application of SXT to whole-volume 3D imaging of multicellular filamentous bacteria remains unexplored. Cable bacteria therefore provide an ideal system to evaluate the ability of SXT to bridge molecular ultrastructure to whole-cell organization in an intact multicellular bacterium.

Here, we apply SXT to map the structural and functional architecture of multicellular *Ca.* Electrothrix filaments. Through an integrated approach that combines SXT, scanning transmission electron microscopy coupled with energy-dispersive X-ray spectroscopy (STEM-EDX), cryo-ET, and fluorescence microscopy, we place previously described ultrastructural features into their whole-cell context and reveal novel aspects of cable bacteria organization. Our volumetric SXT reconstructions reveal extensive structural heterogeneity and dynamic cell-cycle-dependent spatial reorganization of intracellular structures. Furthermore, by integrating mesoscale SXT with cryo-ET and STEM-EDX, we relate differences in X-ray absorption and local biomolecular density to the molecular architecture of the conductive network, distinguishing the junction lamella from the conductive fibers. Finally, exploiting the large 3D field of view, we identify a previously undescribed asymmetric mode of cell division in cable bacteria, providing a framework for how cytokinesis may be coordinated with the maintenance of a contiguous conductive network. Taken collectively, our results establish SXT as a powerful approach for studying multicellular bacterial organization and provide multiscale insights into the structural basis of electrical function in cable bacteria.

## RESULTS

### Whole-cell SXT reveals structural heterogeneity among cable bacteria filaments

To investigate the structural organization of cable bacteria at the whole-cell level, we performed SXT on four intact *Ca.* Electrothrix filaments isolated from a single environmental sediment. This dataset comprised 193 individual cells from four distinct filaments, corresponding to approximately 772 µm of reconstructed filament length. While all filaments displayed the characteristic multicellular organization typical of cable bacteria ^1,6,9^, comparison between filaments revealed pronounced morphological and physiological heterogeneity. Segmentation of intracellular structures based on LAC contrast revealed distinct patterns of nucleoid organization across filaments. Filament 1 exhibited a condensed, spatially confined nucleoid distribution with clear cytoplasmic separation (**Fig. 1A, left**), whereas filaments 2–4 displayed a more dispersed DNA distribution (**Fig. 1A**). Because the macroscopic length of cable bacteria greatly exceeds the SXT field of view, intracellular organization initially appeared uniform within the localized regions analyzed (typically spanning 4–5 cells). However, constructing a large-scale composite image along an extended region of filament 2 revealed variation in both DNA distribution and vesicle organization between different regions of the same filament (**Fig. S1**). This within-filament variation suggests that intracellular organization is not necessarily uniform along a cable bacterium filament and may reflect heterogeneity in cellular physiological or cell-cycle state. Beyond variation in DNA organization and intracellular inclusions, filament diameter also varied among the analyzed filaments. To avoid projection artifacts caused by filament orientation in the longitudinal (XY) plane, filament diameters were measured from circular cross-sections in two-dimensional (2D) XZ orthoslices obtained through full-rotation tomography. Based on measurements from nine distinct cell-cell junctions per filament, filaments 1-3 measured approximately 4 µm in diameter, while filament 4 reached approximately 6 µm (**Fig 1B**).

**Figure 1.**
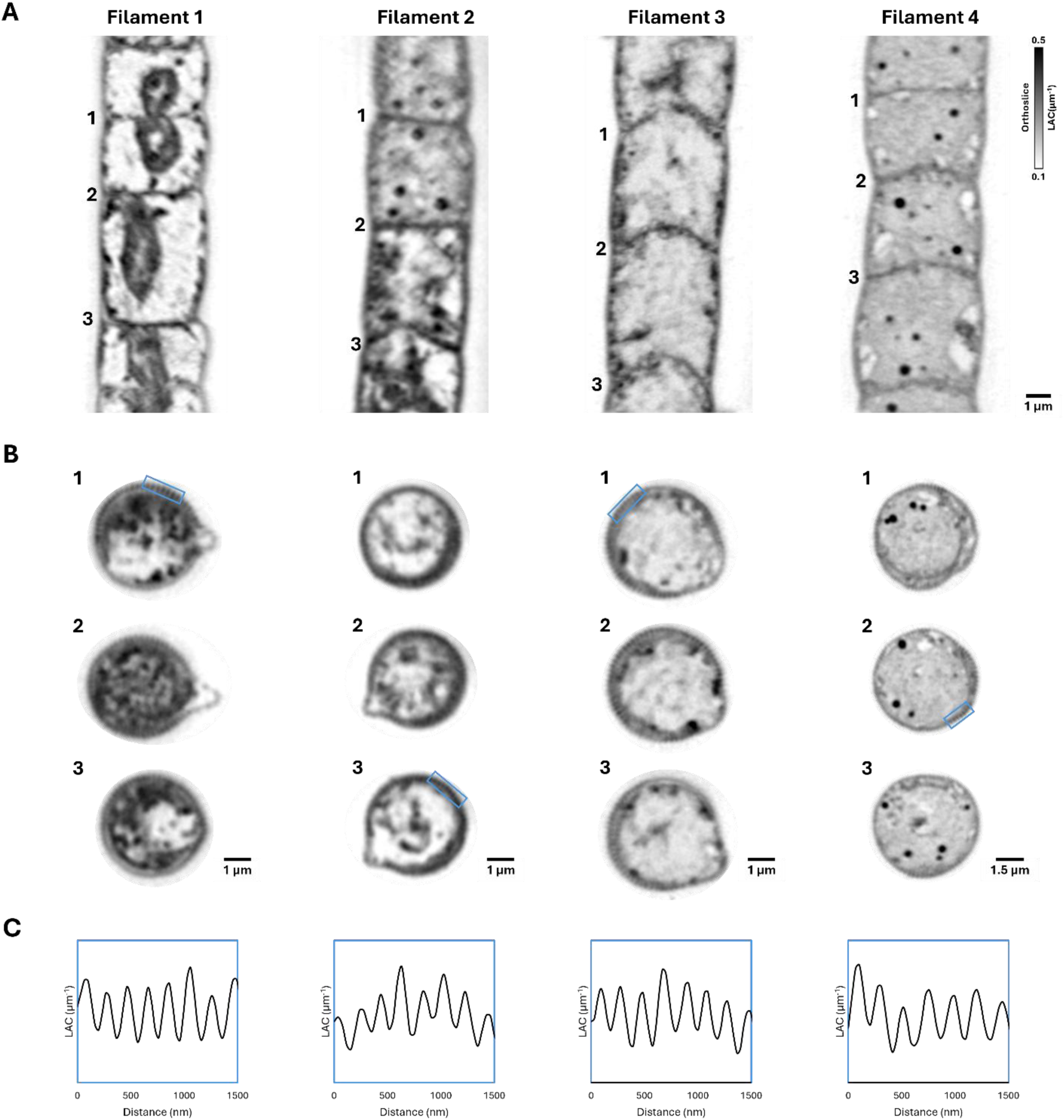
Mesoscale structural heterogeneity, dimensional morphotypes, and periplasmic fiber periodicity in cable bacteria filaments. (A) Representative longitudinal XY SXT orthoslices of four intact cable bacteria filaments isolated from the same sediment sample, illustrating variations in DNA organization and intracellular inclusions. Numbers (1–3) indicate the specific cell–cell junctions of the corresponding transverse XZ orthoslices shown in (B). (B) Cross-sectional 2D XZ orthoslices at three positions along each filament, showing differences in filament diameter and the periodic periplasmic fibers around the cell envelope. (C) Representative linear intensity profiles extracted from the 1.5 µm peripheral regions (blue boxes in B) showing the regular spatial periodicity of the periplasmic conductive fibers across all four filaments.

To assess whether this dimensional and morphological heterogeneity extended to the periplasmic conductive network, we analyzed cross-sectional tomographic slices of multiple cell junctions, where the periplasmic conductive fibers were most clearly visualized as periodic circumferential structures within the cell envelope. LAC profiles extracted from representative 1.5 µm regions of the filament periphery captured the high-resolution periodicity of the conductive fibers **(Fig. 1C).** Fast Fourier Transform (FFT) analysis of the unwrapped annular cell envelope intensity profile identified the dominant periodicity of the conductive fibers, allowing calculation of the total number of parallel periplasmic conductive fibers for each filament **(Fig. S2)**.

The abovementioned analyses identified two structural morphotypes within the same sediment sample: a narrow (4 µm) morphotype (filaments 1-3) containing approximately 60 periplasmic fibers and a wider (6 µm) morphotype (filament 4) with approximately 80 fibers. Both compact and dispersed DNA organizations were observed within each morphotype, suggesting that intracellular organization varies independently of filament diameter and fiber number.

### Structural and chemical organization of the periplasmic fiber apparatus

SXT reconstructions revealed a distinct density contrast between the periplasmic fibers and their junction lamellae at the cell-cell junctions. Specifically, the LAC value of the periplasmic fibers was higher than that of the junction lamellae (**Fig. 2**). The higher LAC of the fibers indicates greater X-ray absorption, consistent with a higher local density of carbon- and nitrogen-rich biomolecules relative to the junction lamellae. Despite differences in fiber count between the two morphotypes, this LAC contrast between the longitudinal periplasmic fibers and junction lamella was consistently observed across all analyzed filaments.

**Figure 2.**
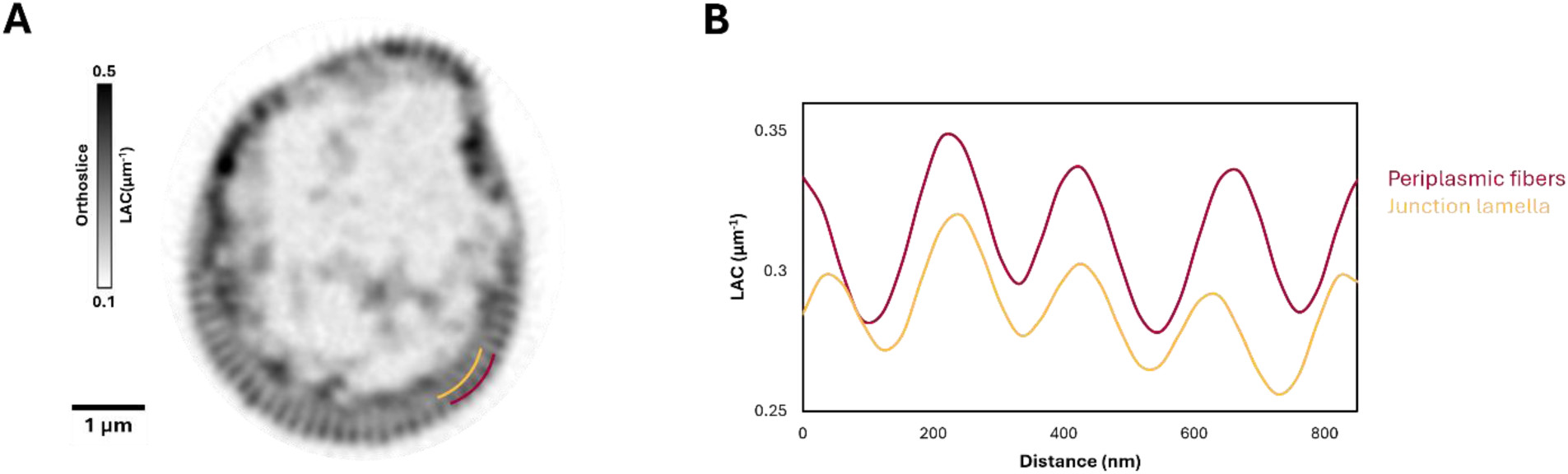
SXT resolves differences in molecular density between periplasmic fibers and the junction lamella. (A) Representative transverse XZ orthoslice at a cell–cell junction used for LAC measurements. (B) LAC profiles measured from the peripheral regions highlighted in (A). Colored lines indicate representative regions corresponding to the periplasmic fibers (magenta) and junction lamella (yellow).

Given the presence of a nickel-containing cofactor in the conductive fibers ^11^, STEM-EDX elemental mapping was performed on fiber skeletons prepared from independent cable bacteria filaments (**Fig. 3A**). Fiber skeletons were previously shown to retain the periplasmic conductive fiber network after chemical removal of the cell membranes and cytoplasmic content ^6–8^. The nickel signal co-localized with the periplasmic fibers and remained continuous across the cell-cell junctions. These observations indicate that the nickel-enriched component of the conductive pathway remains chemically continuous across cell-cell junctions despite the LAC differences observed by SXT.

**Figure 3.**
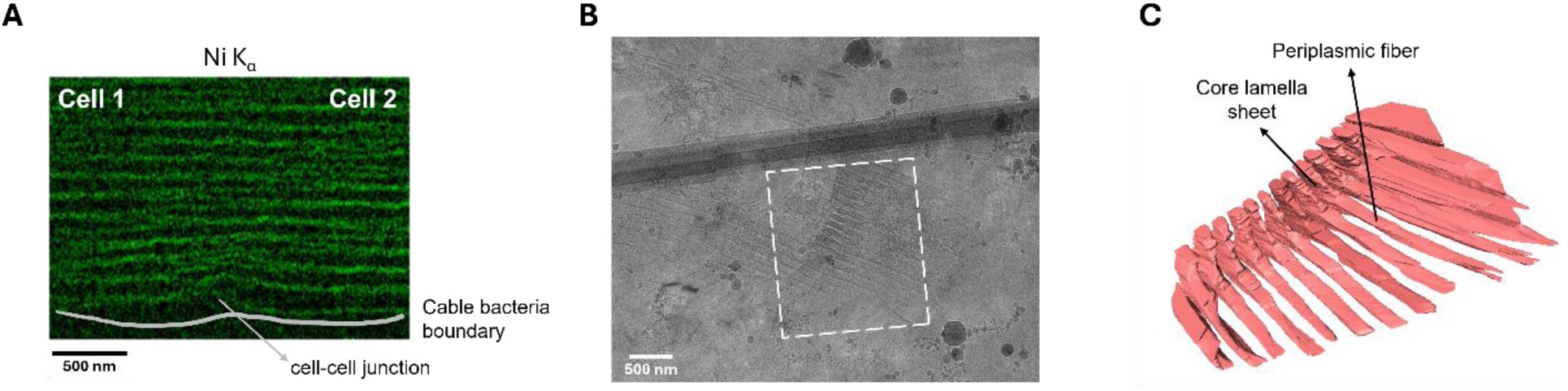
Elemental and ultrastructural organization of the periplasmic fiber network. (A) STEM-EDX elemental map of nickel (Ni Kα) along the fiber skeleton of a cable bacteria filament spanning two adjacent cells. The gray line indicates the outer cable bacteria filament boundary. (B) Representative cryo-ET image of a cryo-FIB-milled lamella*e* from an intact cable bacteria filament. The dashed box indicates the region used for the 3D segmentation shown in (C). (C) 3D segmentation of the cryo-ET reconstruction showing the core lamella sheet and periplasmic fiber*s*.

To complement the structural and elemental information provided by SXT and STEM-EDX, we employed cryo-ET on cryo-FIB-milled lamellae of intact, fully hydrated cable bacteria filaments (**Fig. 3B**). High-resolution cryo-ET reconstructions showed that the core lamella sheets are directly connected to the longitudinal periplasmic fibers (**Fig. 3C**).

### Quantitative mapping of intracellular structures

SXT enables quantitative characterization of intracellular structures through LAC measurements. Because X-ray absorption within the water window depends on the local biochemical composition and biomolecular density, particularly the abundance of carbon- and nitrogen-rich biomolecules, intracellular structures can be distinguished in fully hydrated cells without staining or chemical fixation.

SXT reconstructions identified highly absorbing cytoplasmic inclusions across all analyzed cable bacteria filaments (**Fig. 4A, B**). Complementary fluorescence microscopy on independent cable bacteria filaments isolated from the same environmental sediment, using DAPI to localize DNA and Nile Red to label hydrophobic lipid bilayers, confirmed that these inclusions correspond to lipid-enclosed cytoplasmic vesicles (**Fig. 4C**), consistent with previous ultrastructural observations of membrane-bound vesicles in cable bacteria ^6,9^.

**Figure 4.**
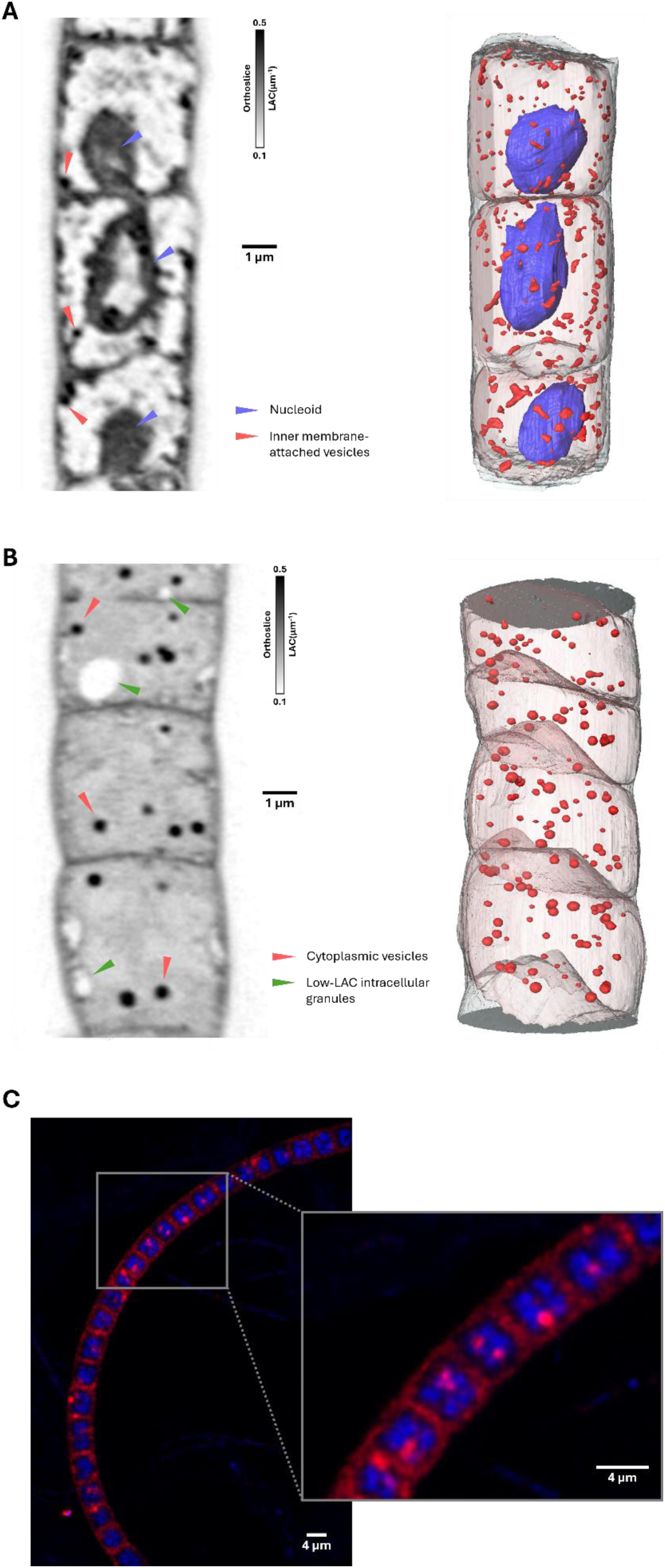
Intracellular organization of lipid-enclosed vesicles and low-LAC granules in cable bacteria filaments. (A) Left: representative longitudinal XY SXT orthoslice of a filament containing actively dividing cells. Red arrowheads indicate inner membrane-attached vesicles, and blue arrowheads indicate the nucleoid. Right: 3D volumetric segmentation of the SXT reconstruction, depicting condensed nucleoids (blue), inner membrane-attached vesicles (red), the inner membrane (transparent pink), and outer membrane (semi-transparent gray). (B) Left: representative longitudinal XY SXT orthoslice of a filament without evident active cell division. Red arrowheads indicate round cytoplasmic vesicles, and green arrowheads indicate low-LAC intracellular granules. Right: 3D volumetric segmentation of the SXT reconstruction, depicting cytoplasmic vesicles (red), the inner membrane (transparent pink), and outer membrane (semi-transparent gray). (C) Representative fluorescence microscopy image of an independent filament co-stained with DAPI (blue, DNA) and Nile Red (red, lipid-rich structures).

Volumetric SXT segmentation and normalized Euclidean distance transform (EDT) mapping identified distinct patterns of vesicle morphology and intracellular localization across filaments (**Fig. 4A, B**). In filaments containing actively dividing cells, vesicles were predominantly irregular in shape and attached to the inner membrane (inner membrane-attached vesicles, IMAVs; **Fig. 4A**; 506 vesicles analyzed across 5 cells, corresponding to a total analyzed cell length of 23.7 µm), with their normalized EDT distribution strongly skewed toward the cell periphery (**Fig. S3A**) and a mean LAC value of 0.45 ± 0.01 µm⁻¹ (**Video S1**). In contrast, filaments without evident cell division contained predominantly round cytoplasmic vesicles (CVs; **Fig. 4B**; 253 vesicles analyzed across 5 cells, corresponding to a total analyzed cell length of 28.6 µm) characterized by a broader intracellular distribution shifted toward the cell interior (**Fig. S3B**) and a comparable mean LAC value of 0.45 ± 0.03 µm⁻¹ (**Video S2**). Both vesicle populations exhibited broad size distributions, with diameters ranging from approximately 50 to 400 nm, although cytoplasmic vesicles showed a distribution shifted toward larger diameters relative to inner membrane-attached vesicles (**Fig. S4**).

In addition to lipid-enclosed vesicles, SXT reconstructions identified a second class of intracellular inclusions characterized by low LAC values (0.13 ± 0.04 µm⁻¹, **Fig. 4B**, green arrow), consistent with a lower density of carbon- and nitrogen-rich biomolecules relative to the surrounding cytoplasm and compositionally distinct from the lipid-enclosed vesicles. STEM-EDX elemental mapping on independent cable bacteria filaments isolated from the same sediment demonstrated that morphologically equivalent granules were strongly depleted in carbon and highly enriched in oxygen. Furthermore, these inclusions were also heavily enriched in phosphorus, which co-localized with magnesium, calcium, and sodium (**Fig. 5**), providing a multi-elemental signature consistent with poly-P granules. Similar to lipid vesicles, the size and distribution of these poly-P granules varied widely both between and within individual filaments (**Fig S5**).

**Figure 5.**
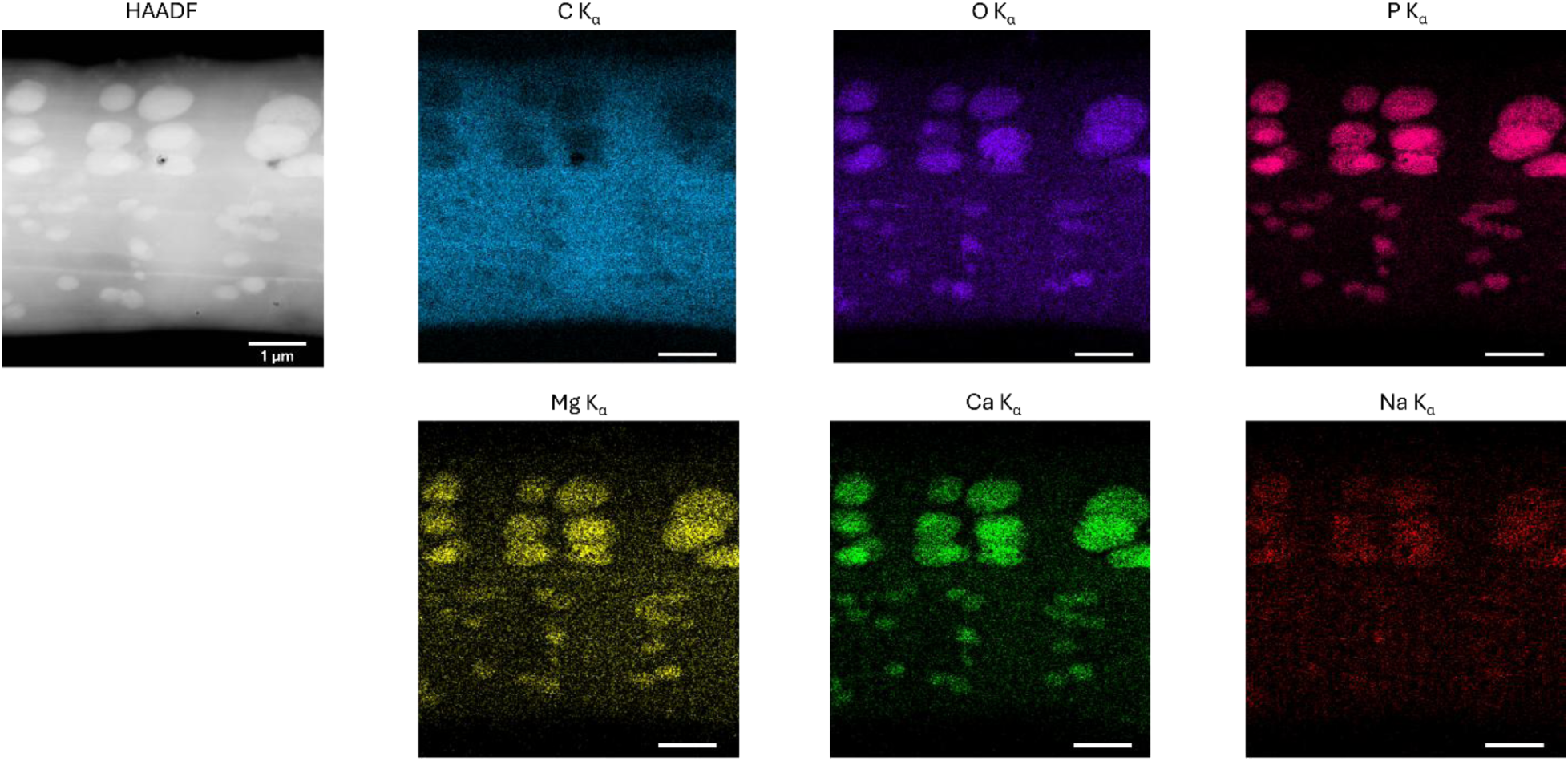
STEM-EDX elemental mapping of poly-P granules in a cable bacteria filament. High-angle annular dark-field (HAADF) image and corresponding energy-dispersive X-ray spectroscopy (EDX) elemental maps (K_α_ lines) of an intact cable bacteria filament showing the spatial distribution of carbon, oxygen, phosphorus, magnesium, calcium, and sodium. Scale bars, 1 µm.

### Nucleoid remodeling and asymmetric cytokinetic constriction

SXT reconstructions identified cells at different stages of the division cycle, characterized by differences in cell length and nucleoid organization. Smaller cells typically contained a single high-LAC domain embedded within the nucleoid, whereas elongated cells consistently exhibited two high-LAC nucleoid-associated domains positioned at opposite poles of the condensed nucleoid (**Fig. 6A**). 3D segmentation and higher-magnification views resolved these nucleoid-associated domains as ring-like structures consisting of a high-LAC shell surrounding a lower-density core (**Fig. 6B, C** and **Video S3**). Representative LAC line profiles across individual domains further supported this organization, showing lower LAC values in the central region relative to the surrounding high-LAC shell (**Fig. S6**). The high-LAC nucleoid-associated domains had an average diameter of approximately 400 nm and occupied, on average, approximately 1% of the corresponding nucleoid volume (n = 6 domains; **Video S4**).

**Figure 6.**
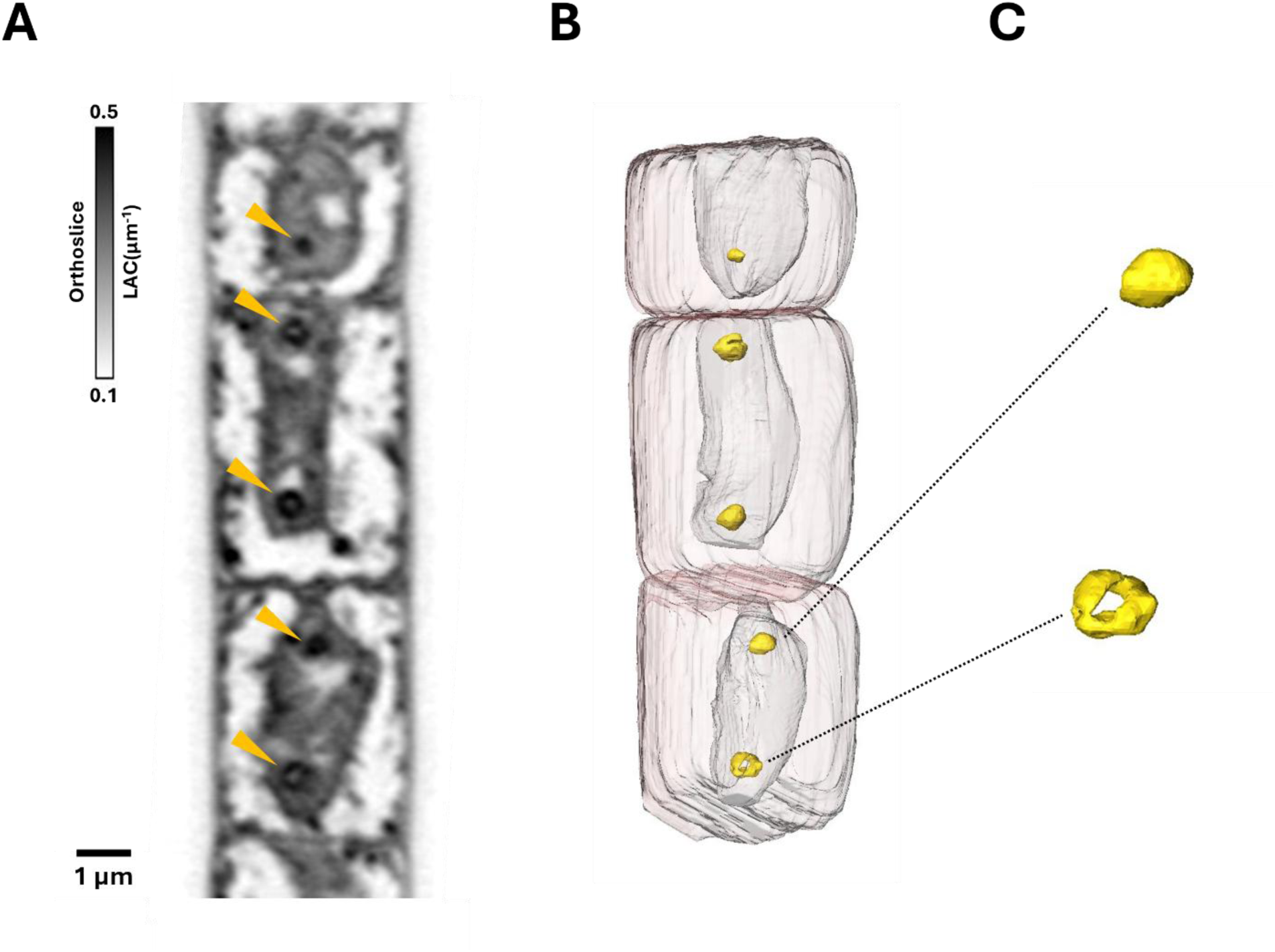
Nucleoid organization and high-LAC nucleoid-associated domains during cell division in cable bacteria filaments. (A) Representative longitudinal XY SXT orthoslice of a dividing cable bacteria filament showing condensed nucleoids containing high-LAC domains (yellow arrowheads). (B) 3D volumetric segmentation of the SXT reconstruction shown in (A), depicting the spatial distribution of the nucleoids (semi-transparent light gray), high-LAC nucleoid-associated domains (yellow), and inner membrane (transparent pink). (C) Magnified 3D renderings of the high-LAC nucleoid-associated domains indicated by the dashed lines in (B).

Further examination of dividing cells unveiled an atypical pattern of asymmetric cell division characterized by unilateral membrane constriction. While non-dividing cell junctions exhibited a symmetric density distribution across the cell-cell junction interface, dividing junctions showed unilateral membrane invagination, with constriction initiating predominantly from one lateral side of the cell (**Fig. 7A, B**). This asymmetric constriction pattern was observed across multiple dividing cells along the same filament.

**Figure 7.**
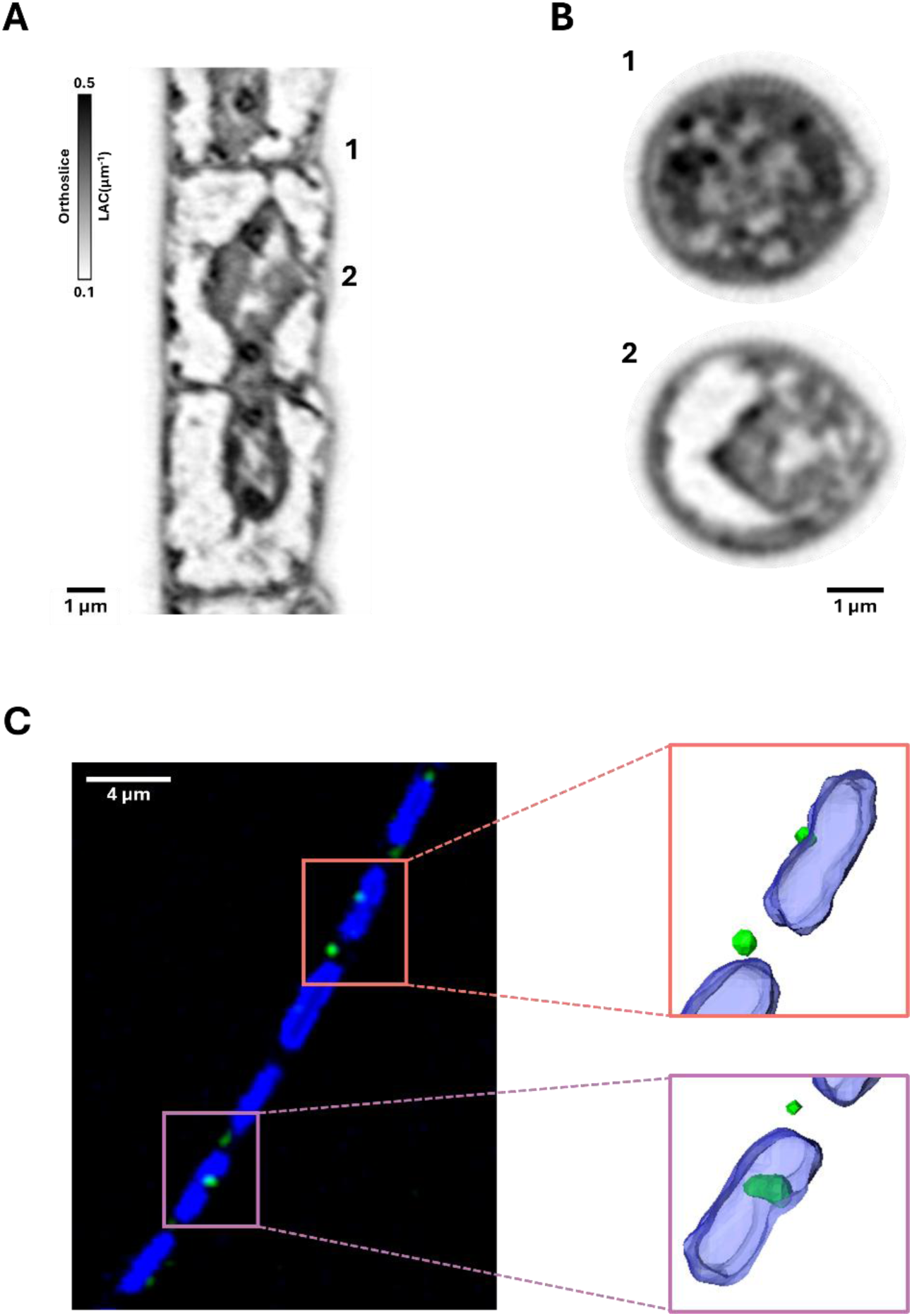
Asymmetric cell constriction and unilateral FtsZ localization during cell division in cable bacteria filaments. (A) Representative longitudinal XY SXT orthoslice of a dividing cable bacteria filament. Numbers 1 and 2 indicate the positions of non-dividing and dividing cell junctions, respectively, corresponding to the transverse orthoslices shown in (B). (B) Cross-sectional 2D XZ orthoslices at the non-dividing cell junction (position 1, top) and the actively dividing cell junction (position 2, bottom) shown in (A). (C) Fluorescence microscopy of an independent dividing filament co-stained with DAPI (blue, DNA) and an FtsZ-specific fluorescent probe (green). Enlarged 3D renderings of the indicated regions show the unilateral spatial localization of FtsZ (green) relative to the DNA (blue).

Fluorescence microscopy targeting the cell division protein FtsZ confirmed that the structural asymmetry observed by SXT was accompanied by asymmetric localization of the division machinery. Co-staining with DAPI to visualize the nucleoid and an FtsZ-specific fluorescent probe showed that FtsZ was not symmetrically distributed across the division plane. Instead, FtsZ was frequently localized to one lateral region of the dividing cell. 3D rendering of the segmented cells further corroborated this pronounced unilateral localization of FtsZ (**Fig. 7C** and **Video S5**).

## DISCUSSION

SXT was previously applied to a limited number of bacterial systems ^25^, and its potential for investigating multicellular bacterial organization has remained unexplored. Here, we demonstrate that SXT provides a mesoscale view of multicellular cable bacteria by generating a volumetric 3D atlas of intact cells in a near-native state. The resulting atlas comprises SXT tomographic reconstructions of 193 cells across four intact filaments, together with associated 3D segmentations and supplementary video reconstructions (**Videos S1–S5**), with the underlying imaging datasets deposited in EMPIAR-47488567.

By complementing SXT with cryo-ET, STEM-EDX, and fluorescence microscopy, we further establish a multiscale imaging framework in which molecular-scale observations can be interpreted in a whole-cell context. SXT provides the volumetric context and quantitative X-ray absorption contrast needed to relate distinct biomolecular features to whole-cell architecture ^25,27,29,31^. This integrative approach places previously described structures, including the periplasmic conductive fibers, junction lamellae and their core lamella sheets, intracellular vesicles, and poly-P granules, into their native 3D organization across multiple cells within a filament. Importantly, this whole-cell perspective also reveals previously unresolved features of cable bacteria organization, including intracellular heterogeneity associated with cell-cycle state, differences in biomolecular density within the conductive apparatus, and asymmetric cytokinetic constriction.

Two distinct morphotypes, distinguished by filament diameter and the number of periplasmic fibers, coexisted within the same sediment sample (**Fig. 1**). The morphotypes were characterized by filament diameters of approximately 4 and 6 µm (**Fig. 1B**). Large filament diameter is a defining morphological characteristic of *Ca.* Electrothrix gigas, with cells wider than approximately 2.5 µm and commonly ranging from 2.5 to 8 µm, approximately 2–10-fold wider than other cable bacteria species ^32,33^. The *Ca.* Electrothrix sp. NPCB-01 recovered from the same sediment sample as the filaments analyzed here is phylogenetically closely related to *Ca.* Electrothrix gigas ^34^. Notably, the approximately 60 and 80 periplasmic fibers resolved in our two morphotypes (**Fig. 1C**) extend the previously reported range of approximately 12–62 fibers ^1,2,6,9,32,35^, with the wider morphotype containing, to our knowledge, the highest fiber count reported in cable bacteria to date. This relationship between filament diameter and conductive fiber number is consistent with previous structural studies showing that the number of conductive fibers scaled with filament diameter ^6^. In contrast, DNA organization and vesicle morphology and distribution varied independently of filament diameter and fiber number, with both compact and dispersed DNA distributions, as well as inner membrane-attached and cytoplasmic vesicles, observed within the same morphotype (**Figs. 1A** and **4**). Together, these observations point to heterogeneity at two levels: variation in filament morphology, reflected by diameter and conductive fiber number, and variation in intracellular organization, including DNA distribution and vesicle morphology, which appears to be associated with cell-cycle and physiological state. Whether the observed morphotypes represent distinct populations or phenotypic variation within natural cable bacteria populations remains unknown. Previous studies have similarly reported substantial variation in filament diameter and fiber number among cable bacteria filaments coexisting in the same sediment enrichment and have suggested that these morphological differences may reflect considerable intraspecific phenotypic variation rather than taxonomic differentiation ^2,6,9,36^.

Previous ultrastructural studies established that the longitudinal conductive fibers and junction lamella are distinct components of the cable bacteria conductive apparatus ^9^. Our SXT measurements reinforce this distinction by showing that these regions exhibit quantitatively different X-ray absorption properties: the junction lamella exhibits lower LAC values than the periplasmic fibers, consistent with a lower local molecular density (**Fig. 2**). This provides, to our knowledge, the first quantitative *in situ* evidence that these two components differ in local biomolecular density within intact cable bacteria filaments. Despite this biochemical differentiation, our high-resolution cryo-ET reconstructions resolved a physical connection between the core lamella sheets and longitudinal fibers (**Fig. 3B, C**), supporting the recently proposed model in which the core lamella sheets function as an interconnection hub linking conductive fibers across cell–cell junctions, thereby maintaining LDET ^9^. This structural continuity was further supported by STEM-EDX, which showed that the nickel-enriched conductive pathway was continuous across the cell–cell junctions (**Fig. 3A**), consistent with previous reports localizing nickel to the conductive fibers ^9,11^. Together, these observations indicate that differences in local biomolecular density are integrated within a physically and chemically continuous conductive network, illustrating how complementary imaging modalities connect mesoscale organization with molecular ultrastructure.

Beyond the conductive network, our volumetric reconstructions demonstrate pronounced variation in intracellular organization associated with cell-cycle state (**Fig. 4**). In filaments exhibiting clear division-associated morphologies, including elongated cells and envelope constrictions, vesicles were predominantly irregular in shape and localized near the inner membrane, whereas filaments without evident cell division contained mainly rounded vesicles distributed within the cytoplasm. Intracellular vesicles have previously been observed in cable bacteria, with inner membrane-attached vesicles proposed to function as hotspots for membrane-dependent cellular processes, potentially including components involved in electron transport ^9^. Our observation that inner membrane-attached vesicles predominate in actively dividing filaments adds a cell-cycle dimension to this model, raising the possibility that the additional membrane surface provided by these structures supports electron transfer and other membrane-associated processes during periods of increased metabolic and biosynthetic demand, while potentially also contributing to the extensive membrane remodeling required during cell growth and cytokinesis ^37^. The comparable LAC values of inner membrane-attached and cytoplasmic vesicles further suggest that their contrasting morphology and localization are not accompanied by major differences in overall biomolecular composition. Whether inner membrane-attached and cytoplasmic vesicles represent functionally distinct vesicle populations or whether their morphology and membrane association change during the cell division cycle remains unknown.

In addition to lipid-enclosed vesicles, poly-P granules represent a chemically distinct intracellular storage system in cable bacteria (**Fig. 5**). Poly-P accumulation has previously been documented in both marine and freshwater cable bacteria, with substantial variation in granule size and abundance across and within filaments ^12–15^. Because cable bacteria continuously reposition themselves across steep and shifting sediment redox gradients, individual cells can transition between growth-supporting conditions in suboxic layers and exposure to oxygen at the oxic interface. Previous work proposed that poly-P in cable bacteria may serve not only as an intracellular reserve of energy and phosphate but also support survival during non-growing stages of the cell cycle and provide protection against oxidative stress during exposure to oxic conditions, where cells continue to perform oxygen reduction despite the absence of a known energy-conserving terminal oxidase ^12^. The presence and distribution of poly-P granules observed in our data do not, by themselves, establish their physiological function, but their pronounced heterogeneity suggests that poly-P accumulation in cable bacteria is associated with cellular physiological state rather than representing a uniform storage pool throughout the filament.

More broadly, the coexistence of distinct intracellular compartments and storage inclusions highlights the importance of spatial organization in supporting the multicellular physiology of cable bacteria. Comparable forms of intracellular organization have been described in other sulfur-oxidizing bacteria. In *Candidatus Thiomargarita magnifica*, DNA and ribosomes are organized within membrane-bound compartments associated with localized protein synthesis, while ATP synthase is distributed along an extensive intracellular membrane network ^38^. In vacuolated *Beggiatoa*-like filaments, large intracellular vacuoles can occupy most of the cellular volume and accumulate high concentrations of nitrate, an organization associated with adaptation to fluctuating environmental conditions ^39^. Although these intracellular compartments differ structurally and functionally from the vesicles and poly-P granules observed in cable bacteria, they illustrate how intracellular organization can support distinct physiological and metabolic requirements in bacteria exposed to heterogeneous environmental conditions. In cable bacteria, this intracellular organization occurs within a multicellular framework that requires coordination and complex interactions among cells experiencing different local redox conditions while remaining metabolically interconnected through LDET ^24^.

Our whole-cell volumetric atlas further provides a structural link between nucleoid architecture and the cell cycle in cable bacteria. Previous SXT studies on model single-celled bacteria have demonstrated that nucleoids are internally heterogeneous, containing spatially distinct regions with different X-ray absorption properties, including high- and low-absorbance nucleoid domains as well as internal core-like regions ^40,41^. Our SXT reconstructions of cable bacteria identified a distinct form of intra-nucleoid organization characterized by ring-like high-LAC domains (**Fig. 6**). Their organization follows a cell-cycle-associated pattern, with smaller cells typically containing a single domain and elongated cells containing two domains positioned at opposite poles of the condensed nucleoid. The transition from one to two domains with cell elongation suggests that these structures are associated with chromosome reorganization during the cable bacteria division cycle, although their relationship to chromosome replication or segregation remains unclear. Furthermore, active division was observed across the entire analyzed length of the filament, consistent with the non-apical mode of growth previously proposed for cable bacteria ^42^, with individual cells within the same filament captured at distinct stages of the cell cycle.

This nucleoid reorganization associated with the cell cycle is accompanied by an asymmetric spatial organization of the division machinery. Previous fluorescence studies reported FtsZ accumulation at cell– cell junctions in cable bacteria, consistent with an FtsZ-based division apparatus ^10^. Our fluorescence data extend these observations by showing that FtsZ frequently localizes to one lateral region of the division plane rather than being uniformly distributed around the cell circumference (**Fig. 7C**). This asymmetric localization is consistent with fluorescence and super-resolution studies showing that FtsZ organization in dividing bacteria can be discontinuous and dynamic rather than forming a uniformly continuous ring ^43–47^. Consistent with this asymmetric FtsZ localization, SXT revealed unilateral envelope invagination at dividing junctions, with early constriction morphologies restricted to one side of the division plane (**Fig. 7A, B**). This pattern is consistent with cryo-ET observations across diverse bacterial species showing that cytokinesis can initiate asymmetrically on one side of the division plane, accompanied by short FtsZ-like filaments, and that a complete FtsZ ring is not required for constriction to begin ^48^. Cornelissen et al. previously proposed that the junction lamella originates through invagination of the outer envelope during cytokinesis ^6^. Our observations extend this model by showing that early membrane invagination is asymmetric, with constriction initially evident on one lateral side of the cell. Together, the asymmetric localization of FtsZ and unilateral envelope constriction support a model of cable bacteria cell division in which septation is initiated asymmetrically, with localized FtsZ assembly and envelope invagination occurring first on one side of the division plane before constriction becomes established around the cell circumference (**Fig. 8**). Although the functional significance of this asymmetric cytokinesis remains unclear, it provides a plausible structural mechanism by which cable bacteria could progressively reorganize their shared envelope while minimizing disruption to the continuous periplasmic conductive network and thereby potentially maintaining LDET during cell division.

**Figure 8.**
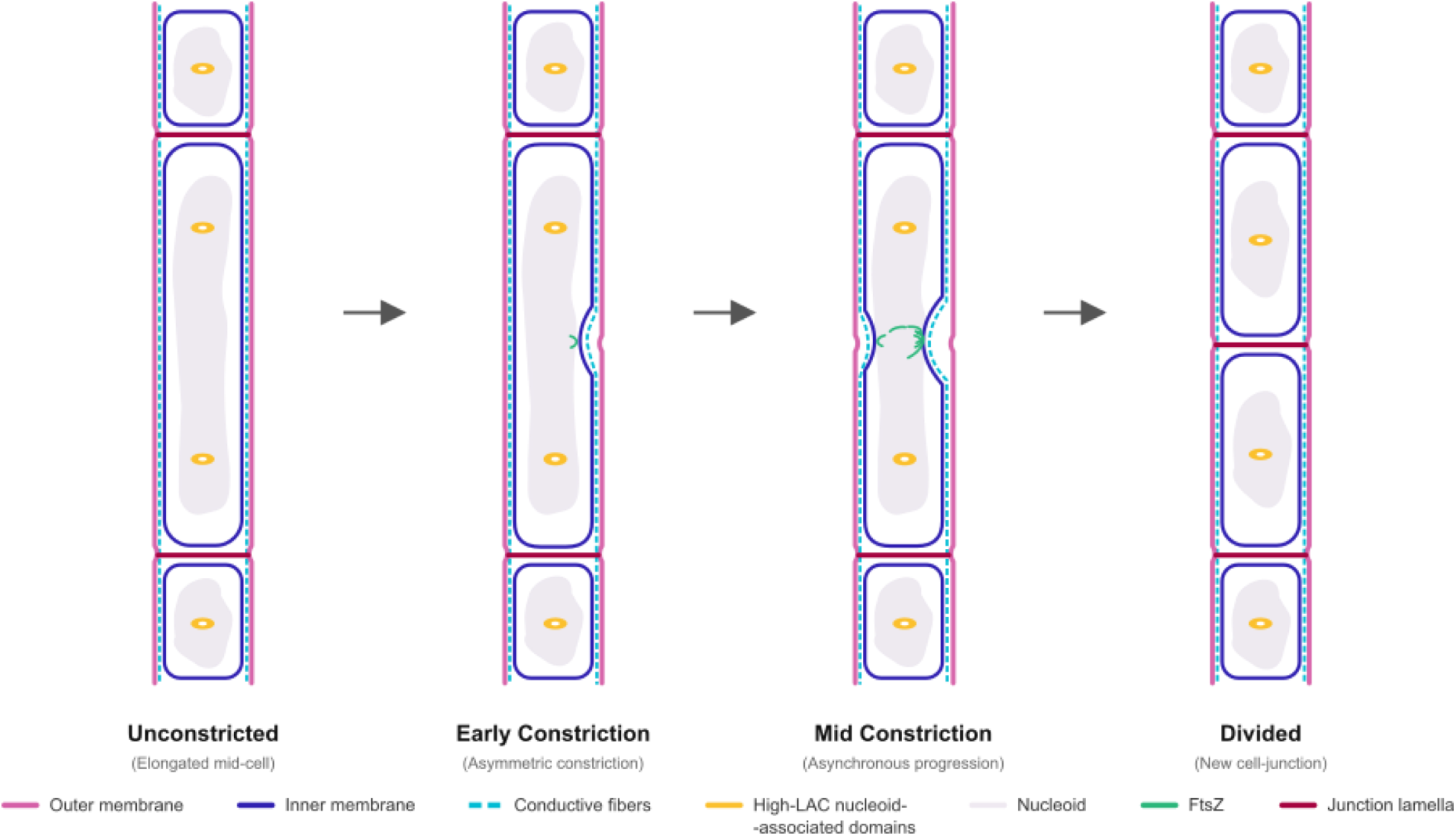
Model of asymmetric cytokinetic constriction in cable bacteria. Schematic diagrams illustrating the progression of cell division based on SXT and fluorescence microscopy observations. In the unconstricted stage, the elongated cell contains two high-LAC nucleoid-associated domains positioned at opposite poles of the condensed nucleoid. Early constriction is characterized by asymmetric FtsZ localization and unilateral membrane invagination at the division plane. As constriction progresses, FtsZ localization and envelope invagination advance asynchronously around the cell circumference, ultimately resulting in the formation of a new cell–cell junction. Periplasmic conductive fibers and junction lamellae are shown to illustrate the organization of the conductive network throughout the proposed division sequence.

## RESOURCE AVAILABILITY

### Lead contact

Requests for further information and resources should be directed to and will be fulfilled by the lead contact, Mohamed El-Naggar.

### Materials availability

This study did not generate new unique reagents.

### Data and code availability

- Original imaging data referenced in the manuscript were submitted to the Electron Microscopy Public Image Archive (EMPIAR). The accession number for the data deposited is 47488567. Data are publicly available as of the date of publication.
- All original code has been deposited at OSF at https://osf.io/dr9aq/overview?view_only=0b00d411da884b3b888aca9e4a59baf5 and is publicly available as of the date of publication.
- Any additional information required to reanalyze the data reported in this paper is available from the lead contact upon request.

## ACKNOWLEDGMENTS

This work was supported by the Gordon and Betty Moore Foundation grant 10148 and the W.M. Keck Foundation award 8626. We thank the California Department of Fish and Wildlife staff for their assistance and sampling permission (permit ID S-222970004-22297-001). We thank the National Center for X-ray Tomography (NCXT) for access to the XM-2 SXT microscope at the Advanced Light Source (LBNL, Berkeley). The NCXT was supported by the NIH (NIGMS P30GM138441) and the Department of Energy’s Office of Biological and Environmental Research Project (DE-AC02-05CH11231). We acknowledge the Core Center of Excellence in Nano Imaging (CNI) at the University of Southern California for support with STEM-EDX imaging and thank Amir Avishai and Lucas Jordao for their assistance with data acquisition. We acknowledge the Midwest Center for Cryo-Electron Tomography (MCCET) at the University of Wisconsin–Madison for cryo-ET data collection. We thank Tingting Yang for insightful discussions and Zenia Motiwala and Cornelius Gati for advice on sample preparation for cryo-ET.

## AUTHOR CONTRIBUTIONS

Conceptualization, A.C. and M.Y.E.-N.; methodology, A.C., V.L., C.A.L., and M.Y.E.-N.; investigation, A.C., A.D., V.L., M.A.M., and M.Y.E.-N.; writing – original draft, A.C. and M.Y.E.-N.; writing – review & editing, A.C., A.D., V.L., M.A.M., C.A.L., and M.Y.E.-N.; funding acquisition, C.A.L., and M.Y.E.-N.; resources, C.A.L., and M.Y.E.-N.; supervision, M.Y.E.-N.

## SUPPLEMENTAL INFORMATION

Document S1. Supplemental Figures S1–S6

Supplemental Videos S1-S5

## STAR★METHODS

### KEY RESOURCES TABLE

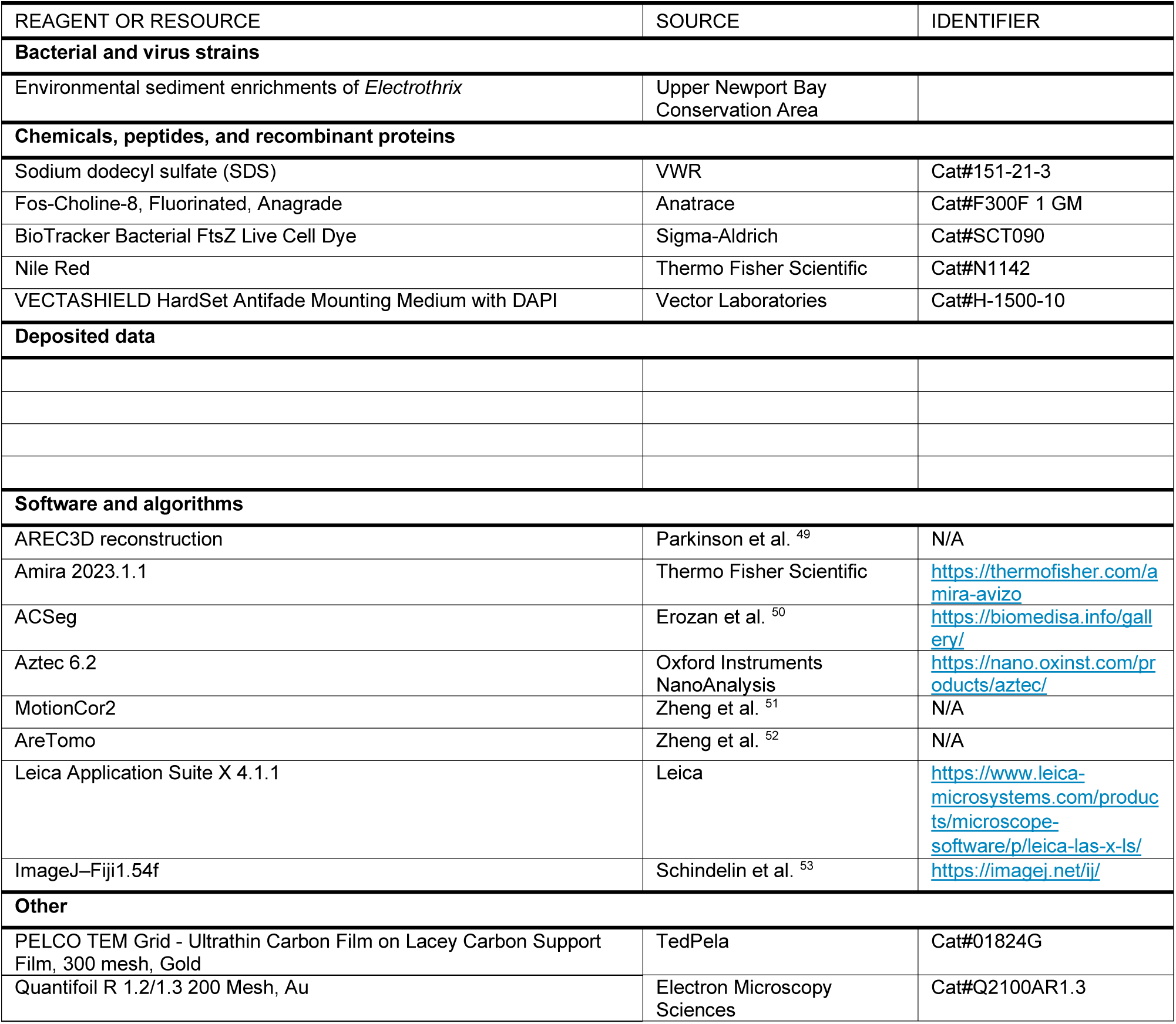

### METHOD DETAILS

#### Cable bacteria incubation

Marine sediment containing cable bacteria was collected from the Upper Newport Bay Conservation Area (33°38’48.0“N, 117°52’14.9“W) ^34^. The top 5-10 cm of sediment was sampled. In the laboratory, the sediment was homogenized and sieved to remove debris and plant material. The processed sediment was subsequently transferred into glass jars, minimizing the presence of trapped air bubbles. The jars were incubated in a dark water bath at 15 °C, fully submerged in circulating, air-saturated 2.6% artificial seawater. After 4-6 weeks of incubation, a dense population of cable bacteria developed in the sediment and was visible upon visual inspection.

#### Cable bacteria filaments pickup

Small sediment cores were collected from the incubation jars using plastic straws (8 mm diameter, 80 mm length) and transferred to small Petri dishes containing 2.6% artificial seawater. Individual cable bacteria filaments were hand-picked under a dissection microscope using glass hooks made from capillary tubes. Filaments were washed at least three times in Milli-Q water droplets to remove sand and debris. The collected filaments exhibited characteristic features of cable bacteria, including multicellular filamentous morphology, and a parallel conductive nanofiber network ^1,6,9^.

#### SXT sample preparation and data collection

Clean cable bacteria filaments suspended in Milli-Q water droplets were loaded into custom-made glass capillaries with diameters ranging from 10 to 12 µm, according to previously established protocols ^30^. Each capillary was then rapidly plunge-frozen in nitrogen-cooled liquid ethane to preserve cellular structure for subsequent X-ray tomography analysis.

SXT was performed using the XM-2 at the National Center for X-ray Tomography, located at Lawrence Berkeley National Laboratory. X-ray projection images were collected at an energy of 517 eV using a 50 nm resolution objective lens. During data collection, the samples were maintained under a steady stream of liquid nitrogen-cooled helium gas to minimize radiation damage. Projection images were sequentially collected around a 180◦ axis of rotation (capillary axis) in 2◦ increments, with an exposure time of 350 ms. Tomograms were reconstructed from the acquired projections using AREC3D, with pixel intensity normalized to attribute accurate LAC values across all cable bacteria filaments ^30,49^.

#### Filament diameter quantification

Filament diameters were quantified directly from reconstructed SXT tomograms. Because measuring filament width in longitudinal (XY) projections can introduce significant dimensional artifacts depending on the tilt angle and spatial orientation of the filament relative to the imaging plane, all diameter measurements were strictly performed on transverse cross-sections (XZ orthoslices). To ensure statistical robustness and account for any potential minor variations along the multicellular structure, diameters were measured at nine independent cell–cell junctions for each analyzed filament.

#### Quantitative analysis of periplasmic fiber periodicity

To automate the count of periplasmic fibers per cell, a custom rotational periodicity algorithm was implemented in Python utilizing the OpenCV, NumPy, and SciPy libraries. Virtual transverse cross-sections (XZ orthoslices) of cell junctions were first processed using Contrast Limited Adaptive Histogram Equalization (CLAHE) to enhance local contrast and resolve the periplasmic fibers. To linearize the roughly circular geometry of the cell envelope, an annular region of interest encompassing the fibers was defined by a central coordinate and a user-defined radial bandwidth. This annulus was mathematically unwrapped into a 2D rectangular strip via polar-to-Cartesian coordinate transformation using bicubic interpolation. The resulting 2D strip was then averaged along its vertical axis to collapse the structural data into a one-dimensional (1D) raw intensity profile, capturing the periodic spatial density of the periplasmic fibers. To isolate the high-frequency structural features from low-frequency background intensity variations, a high-pass filter was applied by subtracting a Gaussian-smoothed background signal from the raw 1D profile. Finally, FFT was applied to the detrended 1D signal. The resulting magnitude spectrum was analyzed to identify the dominant frequency, which corresponds to the expected structural symmetry order (i.e., the total number of fibers across a circular 360° circumference). When cell geometry deviated substantially from a circular cross-section and FFT analysis could not be applied, the number of fibers was counted manually.

#### SXT segmentation and LAC quantification

Segmentation of cable bacteria structures was performed using Amira 2023.1.1 software (Thermo Fisher Scientific). The cell outer and inner membranes were identified based on local LAC contrast relative to the extracellular environment and cytoplasm, respectively. The outer membrane was initially segmented using the ACSeg 3D UNET model on Biomedisa ^50^, followed by manual refinement using the ‘‘paintbrush’’ tool in Amira. The inner membrane and nucleoid were manually segmented by outlining every tenth orthoslice; these were subsequently interpolated to reconstruct the full 3D label field. The inner membrane was characterized by a mean LAC of 0.22 ± 0.07 µm⁻¹. The nucleoid was identified based on its characteristic intracellular localization and distinct LAC contrast, yielding an average LAC value of 0.34 ± 0.06 µm⁻¹. For simplicity, the entire nucleoid was segmented as a single continuous volume. Vesicle segmentation was achieved using a combination of the ‘‘magic wand’’ tool and manual refinement. For individually segmented nucleoids, high-LAC nucleoid-associated domains, and vesicles, mean LAC values were calculated from all voxels within the corresponding 3D label field. Group-level LAC values are reported as the mean ± standard deviation across individual structures.

#### Morphological measurements and vesicles analysis

Morphological parameters, including individual volumes and total counts of vesicles, nucleoids, and high-LAC nucleoid-associated domains, were extracted using Amira software. The volumes were calculated from the number of segmented voxels and converted to equivalent spherical diameters, defined as the diameter of a sphere with the same volume as the segmented structure. To quantitatively evaluate the intracellular position of segmented vesicles relative to the cell boundary, a 3D distance map was generated using the EDT. For each analyzed cell, the EDT map was overlaid onto the volumetric vesicle label field, and the average spatial position of each vesicle was computed. To account for morphological variations in cell shape and size, distance metrics were normalized across the cell by dividing the individual vesicle EDT value by the maximum EDT value (furthest internal point) within that respective cell. Consequently, this generated a standardized spatial scale where a normalized EDT value of 0 corresponds directly to the cell membrane, and an EDT value of 1 represents the deepest cytoplasmic position. Binned frequency distributions of these normalized values were then plotted to quantitatively differentiate membrane-associated from central cytoplasmic localization.

#### STEM-EDX sample preparation and data collection

STEM-EDX spectra of intact cable bacteria filaments and fiber skeletons were recorded on a JEOL JEM2100F transmission electron microscope equipped with an Oxford X-MaxN 100 TLE Windowless Silicon drift detector (SDD). Clean cable bacteria filaments suspended in Milli-Q water droplets were loaded onto a PELCO® ultrathin carbon film supported by a lacey carbon film on a 300-mesh gold grid. For cable bacteria fiber skeletons, 4 μL of 1% (w/v) sodium dodecyl sulfate (SDS) was added to completely cover the grid surface. The SDS-treated grids were then placed in a humid chamber and incubated for 6 hours. After SDS incubation, the samples were washed three times with Milli-Q water and allowed to dry. Individual spectra and element distribution maps were acquired within 3,000,000 and 110,000,000 total counts at 20,000–60,000 times magnification with a beam energy of 200 keV. Aztec software (Oxford Instruments NanoAnalysis) was used to collect and analyze the raw data.

#### Cryo-ET sample preparation and data collection

Clean cable bacteria filaments suspended in 0.02% Fos-Choline-8, Fluorinated were applied to freshly glow-discharged QuantiFoil Au 1.2/1.3, 200 mesh grids, blotted for 4.0–4.5 s, and plunged into liquid ethane using an EM GP2 (Leica Microsystems). Clipped grids were transferred into a dual-beam (SEM/FIB) Aquilos 2 cryo-FIB microscope (Thermo Fisher Scientific) operating under cryogenic conditions. To improve sample conductivity and reduce curtaining artifacts during FIB milling, the grids were first sputter-coated with platinum (20 mA, 25 s), followed by deposition of an approximately 500 nm-thick organometallic platinum layer using the in-chamber gas injection system (GIS). A second platinum sputter coating (20 mA, 7 s) was then applied prior to milling. Lamellae were prepared using AutoTEM Cryo 2.4 (Thermo Fisher Scientific) and milled to a final thickness of approximately 200 nm. After cryo-FIB milling, the clipped frozen grids were imaged using a Titan Krios G3 (Thermo Fisher Scientific) at 300 kV without the fringe-free optical state. Images were acquired on a Gatan BioQuantum GIF-K3 camera (Gatan) in EFTEM mode using a 20-eV slit. The exposure dose was set to 2.89 e–/Å² per projection, resulting in a total accumulated dose of approximately 107 e–/Å² for data collection on lamellae at a magnification of 26,000×. The estimated pixel size was 3.347 Å, with target defocus set to −4.5 μm. Tilt series of cable bacteria lamellae were collected dose-symmetrically from −60° to +48° with 3° increments. Motion correction was performed using MotionCor2 ^51^, followed by tilt series alignment and tomographic reconstruction with AreTomo ^52^. No CTF deconvolution was applied to the reconstructed tomograms. The periplasmic conductive fibers were manually segmented using Amira 2023.1.1 software.

#### Confocal fluorescence imaging

Clean cable bacteria filaments were stained with Nile Red (Thermo Fisher Scientific; excitation: 552 nm; emission: 636 nm), and BioTracker Bacterial FtsZ Cell Dye (Sigma-Aldrich; excitation: 488 nm; emission: 510 nm) ^54^. Cable bacteria filaments were suspended in VECTASHIELD Hardset Antifade Mounting Medium with DAPI (Vector Laboratories; excitation: 341 nm; emission: 452 nm), covered with a coverslip, and sealed with nail polish. The stained cable bacteria filaments were then imaged using a Leica MICA confocal microscope equipped with a HC PL APO CS2 63×/1.20 water immersion objective. Fluorescence images were processed in LASX software (Leica Application Suite X) using Lightning deconvolution. Image z-stacks were then transferred to ImageJ, where maximum-intensity projections were generated. Segmentation of the FtsZ fluorescence signal was performed using the ‘‘magic wand’’ tool in Amira 2023.1.1 software.

