## Supplemental Information for "A 3D Soft X-Ray Structural Atlas of Multicellular Cable Bacteria"

[1]Department of Physics and Astronomy, University of Southern California, Los Angeles, CA, USA
[2]Department of Chemistry, University of Southern California, Los Angeles, USA
[3]Present address: Department of Biophysics and Howard Hughes Medical Institute, University of Texas Southwestern Medical Center, Dallas, TX, USA
[4]Department of Anatomy, School of Medicine, University of California, San Francisco, San Francisco, CA, USA
[5]Molecular Biophysics and Integrated Bioimaging Division, Lawrence Berkeley National Laboratory, Berkeley, CA, USA
[6]Present address: Beamline B24, Diamond Light Source, Harwell Science and Innovation Campus, Didcot, Oxfordshire OX11 0DE, UK
[7]Present address: Division of Geological and Planetary Sciences, California Institute of Technology, Pasadena, CA, USA
[8]Department of Biological Sciences, University of Southern California, Los Angeles, California, USA
[9]Lead contact

**Table S1. Linear absorption coefficient (LAC) values of intracellular structures identified by SXT**
LAC values are reported as mean ± standard deviation. *n* represents the number of individual structures analyzed for each category.

| Structure | n | LAC ($\mu m^{-1}$) |
|---|---|---|
| Nucleoid | 8 | 0.34 ± 0.06 |
| High-LAC nucleoid-associated domains | 13 | 0.48 ± 0.01 |
| Inner membrane-attached vesicles | 506 | 0.45 ± 0.01 |
| Cytoplasmic vesicles | 253 | 0.45 ± 0.03 |

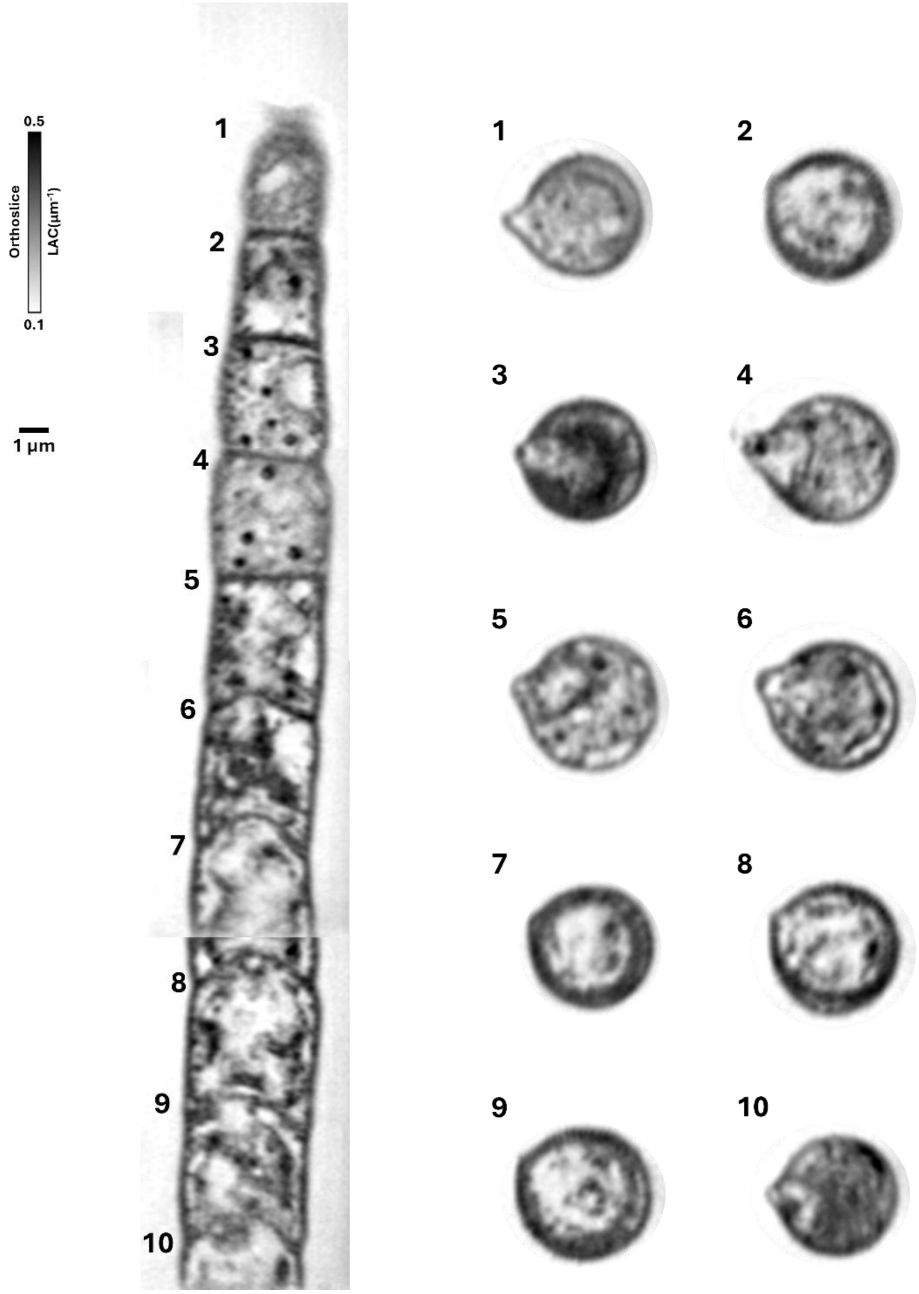


**Figure S1. Longitudinal transition in cellular organization along an extended cable bacteria filament and cross-sectional diameter validation**

Composite longitudinal SXT view of an extended region of filament 2 assembled from consecutive fields of view, showing changes in intracellular organization along the filament. Numbers 1–10 indicate positions along the filament from which representative transverse XZ orthoslices were extracted. The corresponding cross-sections show that filament diameter remains approximately constant along the analyzed region despite apparent variations in width in the longitudinal XY composite.

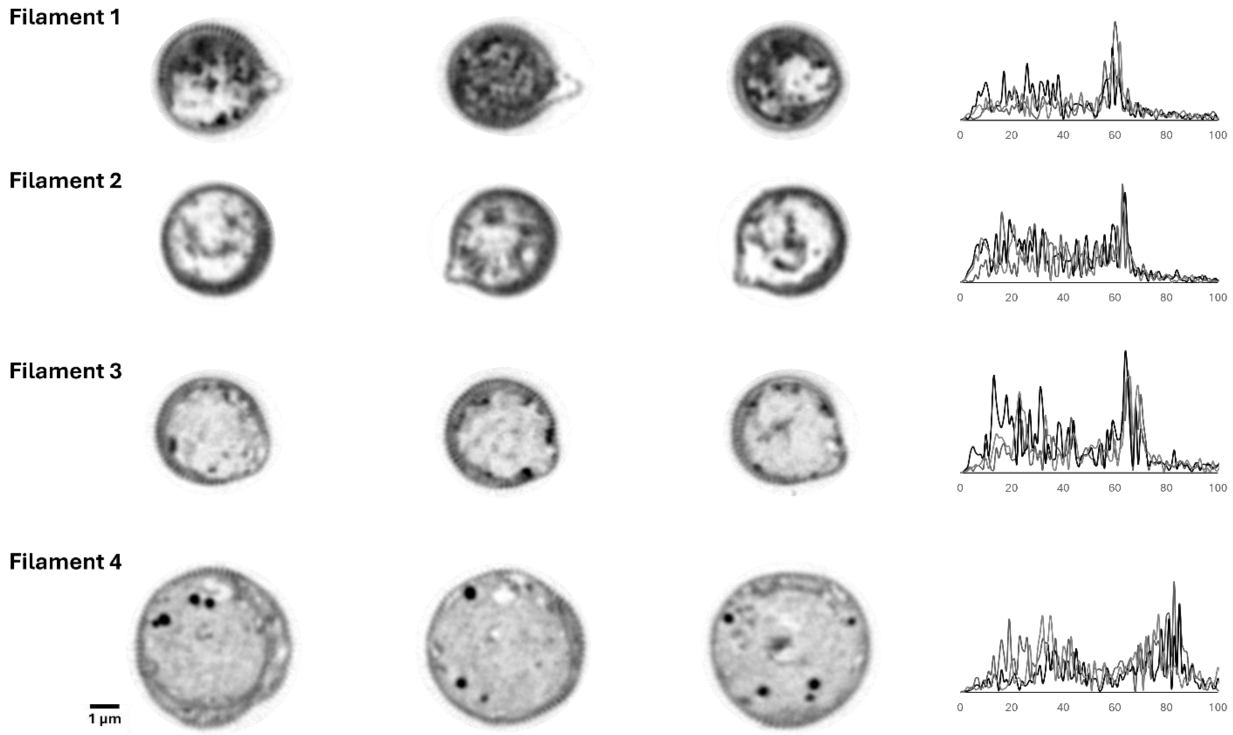


**Figure S2. Quantification of periplasmic fiber numbers across cable bacteria filaments by rotational periodicity analysis**

Representative transverse XZ orthoslices from three distinct cell–cell junctions for filaments 1–4. For each cross-section, the cell envelope was computationally unwrapped to generate a one-dimensional intensity profile, which was analyzed by Fast Fourier Transform (FFT). FFT profiles for the three cross-sections from each filament are overlaid in the corresponding plots on the right. The dominant frequency corresponds to the expected structural symmetry order and was used to estimate the total number of periplasmic fibers across the 360° filament circumference.

A

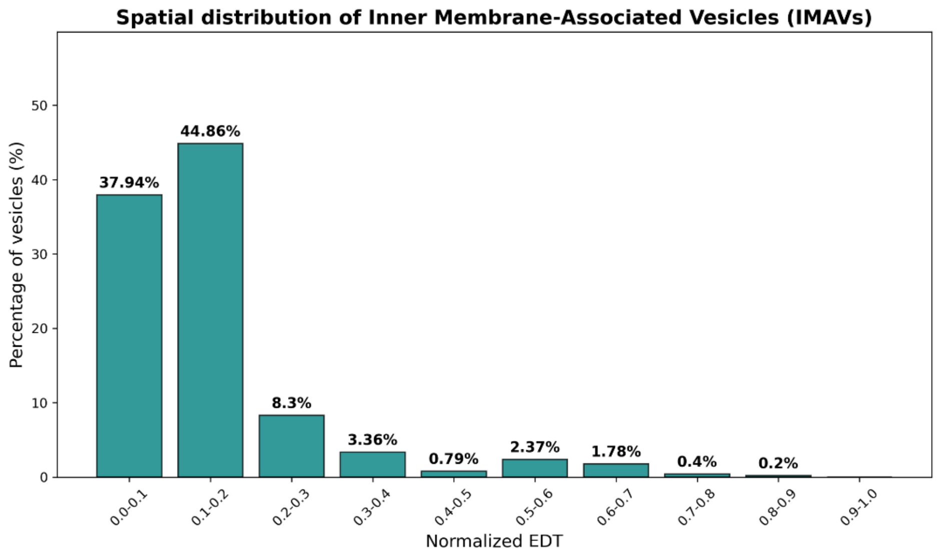


B

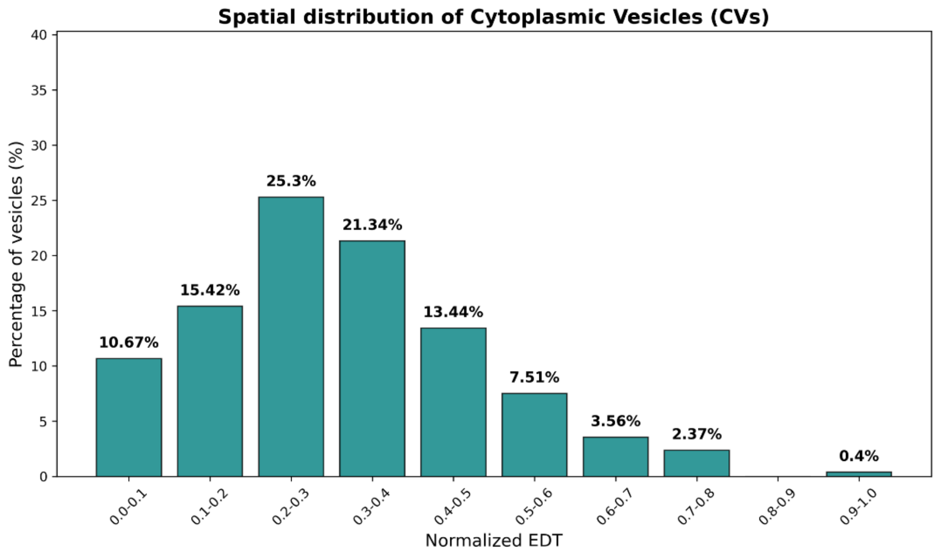


**Figure S3. Quantitative spatial distribution of intracellular vesicles based on normalized Euclidean Distance Transform (EDT)**

Bar charts illustrate the frequency distribution of segmented vesicles across normalized EDT intervals.

(A) In highly active, dividing filaments, irregularly shaped inner membrane-associated vesicles (IMAVs) demonstrate an EDT distribution skewed toward 0, confirming their tight spatial association with the cell envelope.

(B) In less active filaments, round cytoplasmic vesicles (CVs) display a broader spatial distribution with higher EDT values, reflecting a deeper cytoplasmic localization. A normalized EDT value of 0 represents the physical cell membrane, while a value of 1 corresponds to the innermost cellular coordinate furthest from the membrane.

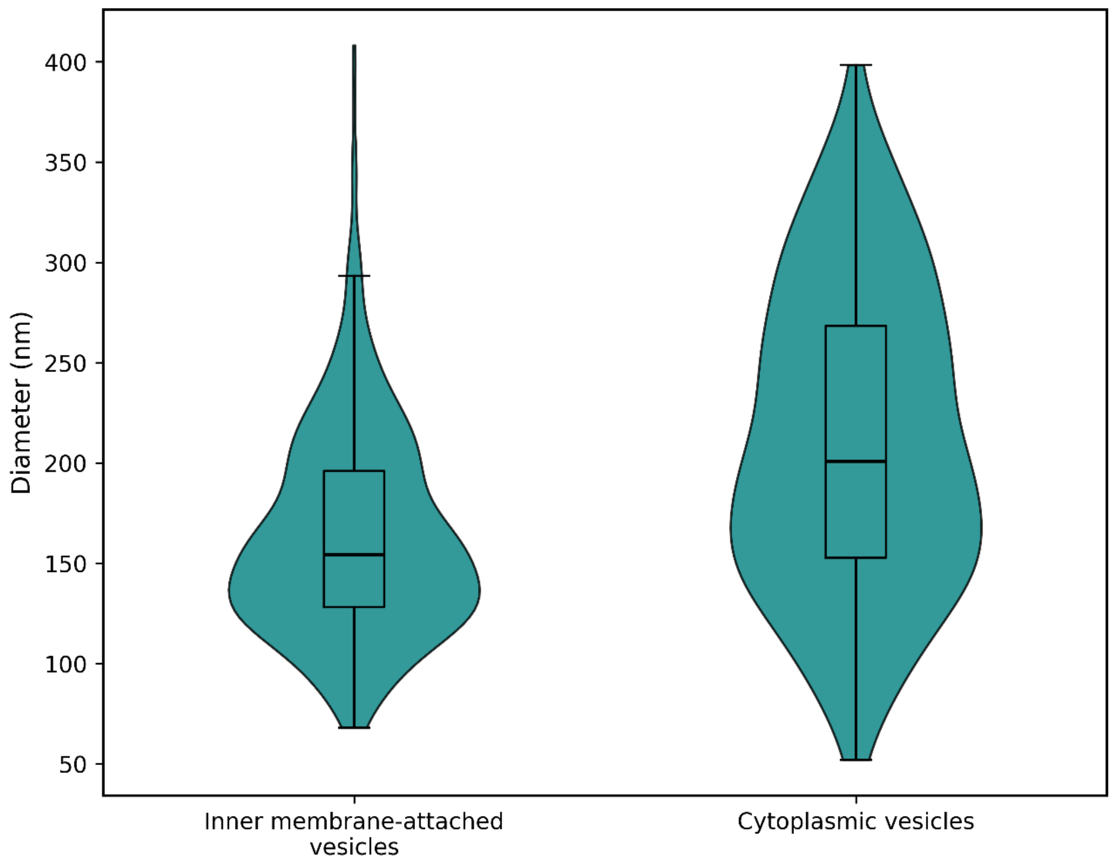


**Figure S4. Size distribution of intracellular vesicles in cable bacteria filaments**
Violin plots showing the distribution of vesicle diameters for inner membrane-attached vesicles (506 vesicles across 5 cells) and cytoplasmic vesicles (253 vesicles across 5 cells). Vesicle diameters were calculated as equivalent spherical diameters from segmented vesicle volumes. Embedded box plots indicate the median and interquartile range.

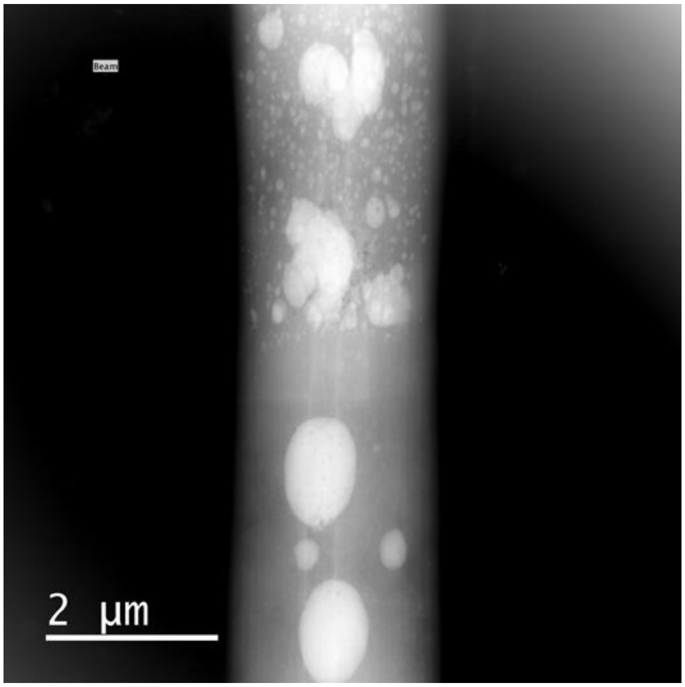


**Figure S5. Heterogeneity in the size and distribution of polyphosphate granules along a cable bacteria filament**

High-angle annular dark-field scanning transmission electron microscopy (HAADF-STEM) image of an intact cable bacteria filament showing intracellular polyphosphate (poly-P) granules of variable size and spatial distribution.

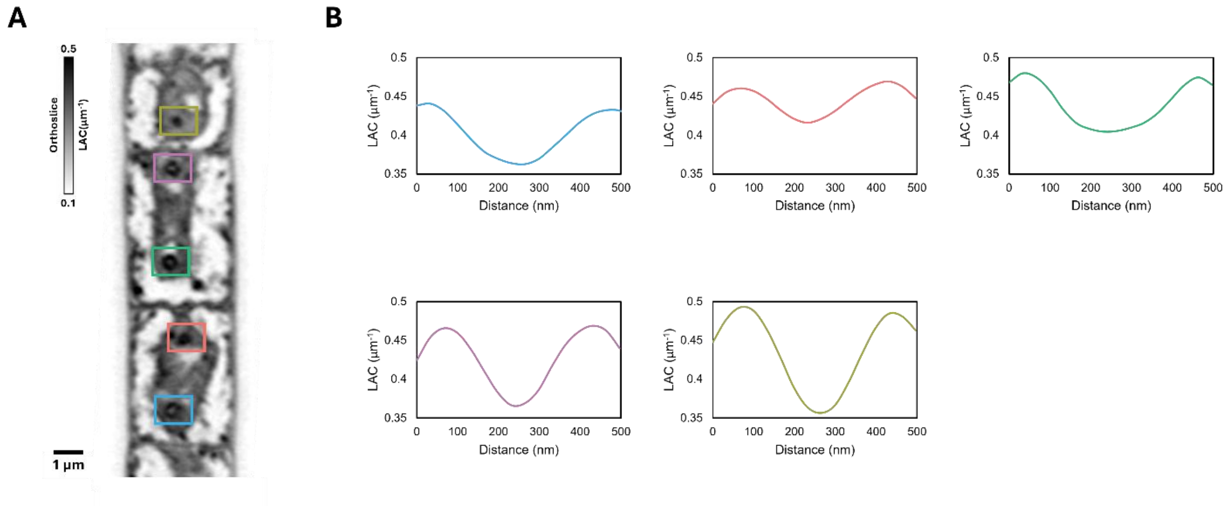


**Figure S6. LAC profiles across high-LAC nucleoid-associated domains**

(A) Representative SXT orthoslice showing five high-LAC nucleoid-associated domains within condensed nucleoids. Colored boxes indicate the regions corresponding to the LAC profiles shown in (B), with matching colors used to identify each domain.

(B) Line profiles across the domains show two regions of elevated LAC separated by a lower-LAC central region, consistent with a ring-like high-LAC shell surrounding a lower-density core.
